# A novel iPSC-based model of ICF syndrome subtype 2 recapitulates the molecular phenotype of ZBTB24 deficiency

**DOI:** 10.1101/2024.04.19.590250

**Authors:** Vincenzo Lullo, Francesco Cecere, Saveria Batti, Sara Allegretti, Barbara Morone, Salvatore Fioriniello, Laura Pisapia, Rita Genesio, Floriana Della Ragione, Giuliana Giardino, Claudio Pignata, Andrea Riccio, Maria R. Matarazzo, Maria Strazzullo

## Abstract

Immunodeficiency, Centromeric instability and Facial anomalies (ICF) syndrome is a rare genetic disorder characterized by variable immunodeficiency. More than half of the affected individuals show mild to severe intellectual disability at early onset. This disorder is genetically heterogeneous and *ZBTB24* is the causative gene of the subtype 2, accounting for about 30% of the ICF cases. ZBTB24 is a multifaceted transcription factor belonging to the Zinc-finger and BTB domain-containing protein family, which are key regulators of developmental processes. Aberrant DNA methylation is the main molecular hallmark of ICF syndrome. The functional link between ZBTB24 deficiency and DNA methylation errors is still elusive. Here, we generated a novel ICF2 disease model by deriving induced pluripotent stem cells (iPSCs) from peripheral CD34^+^-blood cells of a patient homozygous for the p.Cys408Gly mutation, the most frequent missense mutation in ICF2 patients and which is associated with a broad clinical spectrum. The mutation affects a conserved cysteine of the ZBTB24 zinc-finger domain, perturbing its function as transcriptional activator.

ICF2-iPSCs recapitulate the methylation defects associated with ZBTB24 deficiency, including centromeric hypomethylation. We validated that the mutated ZBTB24 protein loses its ability to directly activate expression of *CDCA7* and other target genes in the patient-derived iPSCs. Upon hematopoietic differentiation, ICF2-iPSCs showed decreased vitality and a lower percentage of CD34^+^/CD43^+^/CD45^+^ progenitors. Overall, the ICF2-iPSC model is highly relevant to explore the role of ZBTB24 in DNA methylation homeostasis and provides a tool to investigate the early molecular events linking ZBTB24 deficiency to the ICF2 clinical phenotype.

## 1 Introduction

Immunodeficiency, Centromeric and Facial anomalies syndrome is a rare autosomal recessive disease (1). It is a clinically heterogeneous disorder consisting of combined variable immunodeficiency and variable neurological impairments (2,3). The disease was primarily described as a humoral syndrome but the broad altered clinical spectrum demonstrates a dysregulation of T-cell function (4).

DNA hypomethylation of pericentromeric satellites of chromosomes 1, 9 and 16 is the major molecular hallmark of ICF syndrome, and the basis for the chromosomal instability that is a principal cytological signature of this disease (5,6). ICF syndrome is also genetically heterogeneous. Four subtypes have been described, which are associated with mutations in specific causative genes: the DNA methyltransferase 3B gene (*DNMT3B*), the Zinc-finger and Broad-Complex, Tramtrack and Bric-a-brac domain-containing protein 24 (*ZBTB24*), the cell division cycle-associated protein 7 (*CDCA7*), and the Helicase lymphoid-specific (*HELLS*), for the ICF1-4 subtypes, respectively. No causative gene was identified for the few remaining patients (ICFX)((7–9). There is evidence that these genes contribute to generation and/or maintenance of DNA methylation profiles by interacting with each other in a coordinated manner. However, the precise molecular mechanism underlying these processes is still unclear (10–13). While a subset of genomic regions displays hypomethylation in all disease subtypes, the whole genome methylation signature can distinguish the DNMT3B-mutated ICF1 patients from those of the ICF2-4 subtypes. Moreover, hypomethylation of alpha-satellite is specific to the ICF2-4 patients, while that of pericentromeric satellite repeats is common to all ICF subtypes (10).

*DNMT3B* and *ZBTB24* are the most frequently mutated genes in ICF patients. Subtype 2 accounts for about 30% of ICF cases. In these patients immunodeficiency signs often manifest progressively during their lifetime, characterized as a common variable immunodeficiency (4,14). Autoimmunity predisposition, inflammatory bowel diseases and malignancies have also been reported in ICF2 patients (15,16).

The ZBTB24 transcription factor (TF) belongs to the Zinc-finger (ZF) and BTB domain-containing protein family (17), which includes key regulators of development and differentiation mainly involved in hematopoietic differentiation and T cell function regulation (18). ZBTB24 is less functionally characterized than the other members of the family. It exerts a repressor or activator role by direct binding to DNA targets and through interactions with several partners (19,20). This protein includes three regulatory domains. The N-terminal BTB domain binds co-repressors and histone modification enzymes, contributing to chromatin remodeling (18). This domain also mediates the formation of homo- and hetero-dimers among ZBTB members. The C-terminal C2H2/Krüppel-type Zinc-Finger domain includes eight ZF motifs and mediates the direct binding to specific DNA sequences and other regulatory proteins (20). This domain is responsible for the interaction of ZBTB24 with heterochromatin. ZBTB24 includes a further DNA binding domain known as AT-hook, which binds the minor groove of the DNA double helix and cooperates with the ZF domain for binding to heterochromatin (21). While DNMT3B loss of function (LOF) is clearly associated with DNA hypomethylation, the mechanism/s by which ZBTB24, CDCA7 and HELLS deficiency influence DNA methylation patterns are less obvious and are only beginning to be clarified. Several studies in human and mouse cells demonstrate that ZBTB24 positively regulates the expression of *CDCA7* (*22*). CDCA7 forms a complex with HELLS that functions in nucleosome remodeling (12). The majority of ZBTB24 pathogenic variants are due to nonsense and frameshift mutations causing premature protein termination. The symptoms of patients sharing the same genetic changes can vary both in their clinical manifestations and severity. At least two missense mutations in the ZFs motifs have been observed. They alter the binding ability and the regulatory activity of the protein. The c.1222T>G nucleotide variation is the most frequent missense mutation, reported in eight patients out of 42 described ICF2 cases (15,16). It determines the substitution of a cysteine with a glycine (p.Cys408Gly) in the fifth ZF motif of the protein (23). The clinical phenotype of the patients homozygous for this variant is characterized by a variable spectrum of immunodeficiency and neural disabilities (4,14,16,20,24,25). Notably, three siblings carrying this variant differ in the severity of their immunodeficiency (ranging from a late-onset of severe combined immunodeficiency to asymptomatic reduced level of specific immunoglobulins) and neurological symptoms (4). This demonstrates the complex relationship between the genetic causative mutations and the molecular and clinical phenotypes in chromatin disorders.

ZBTB24 LOF gives rise to early phenotypic alterations. Indeed, mouse models carrying homozygous mutations in the BTB domain show early embryonic lethality (22). Also, a recent study using *Zbtb24* ICF-like mutant murine embryonic stem cells (mESCs) suggested that ZBTB24 is required for the correct *de novo* establishment of DNA methylation through the recruitment of DNMT3B (11). Further disease models are required to investigate the stage at which the epigenetic defect arises. We generated a novel human cellular model of ICF2 that recapitulates the earliest molecular events associated with ZBTB24 deficiency.

## 2 Methods

### 2.1 Generation of iPSCs from CD34^+^ cells

Reprogramming of peripheral CD34^+^-blood cells was conducted using the CD34^+^ Progenitor Reprogramming Kit (StemCell Technologies) according to the manufacturer’s protocols. Briefly, CD34^+^ cells were isolated by positive selection from PBMCs of the ICF2 patient pYM and a healthy donor (control-iPSCs), and expanded with CD34^+^ expansion medium. The Epi5 Episomal iPSC Reprogramming Kit vectors (ThermoFisher) were transfected into the isolated CD34^+^ cells with Nucleofector (Lonza) and Amaxa Human CD34^+^ Cell kit. Cells were transferred to a Geltrex Matrix (ThermoFisher) coated plate in ReproTeSR medium (StemCell Technologies) until rounded colonies appeared. The colonies were isolated and expanded in StemMACS iPS-Brew-XF (Miltenyi). Each clone was passaged with StemPro Accutase (ThermoFisher) and RevitaCell Supplement (ThermoFisher) was added after each passaging. Karyotype analyses were performed on the generated iPSCs following at least twenty passages.

### 2.3 Quantitative Real-Time PCR (qPCR)

RNA derived from iPSCs was reverse transcribed using Random Primers and SuperScript II Reverse Transcriptase (Invitrogen) in a T100 Thermal Cycler (BioRad), according to the manufacturer’s protocol. qPCR was performed using SsoAdvanced universal SYBR Green supermix in a BioRad CFX Opus 96 Real-Time PCR, according to manufacturer’s protocols. Expression levels were normalized to the *GAPDH* gene by the ΔΔCt method. Primer sequences for gene expression are reported in Supplementary Table S1.

### 2.4 Detection of episomal vector by PCR

iPSC suspension was centrifuged, the cell pellet washed with PBS1X and then resuspended in Lysis solution (NaOH 25mM, EDTA 0,2mM). Samples were incubated at 95°C and then treated with Neutralizing solution (TrisHCl 40mM). Cell lysate was used in a PCR experiment with primers specific for the OriP sequence. Primer sequences are reported in Supplementary Table S1.

### 2.5 In vitro differentiation to the three germ layers

iPSCs at 70-80% confluency were treated with Cell Dissociation Buffer (Gibco) to obtain small cell aggregates that were resuspended in KSR medium (Knockout DMEM F12, GlutaMAX 2mM, Knockout SR 20%, NEAA 1mM, β-Mercaptoethanol 0.1mM) and transferred in a 60mm Petri dish with the addition of RevitaCell. Following 7–10 days in suspension culture, embryoid bodies (EBs) were plated in a gelatin-coated 24-well culture dish with coverslips. Immunofluorescence experiments were performed on differentiating cultures utilizing GATA4, βIII-tubulin and MF20 markers to identify endoderm, ectoderm and mesoderm, respectively.

### 2.6 Immunofluorescence staining

iPSCs and EBs were grown on coverslips and fixed with 4% paraformaldehyde and permeabilized with 0.1% Triton X-100. Non-specific binding was blocked with 10% NGS, 0.1% Triton X-100. Primary antibodies against NANOG (SCTB, sc-293121), DNMT3B (Diagenode, pAb-076-005), OCT3/4 (SCTB, sc-5279), GATA4 (SCBT, SC25310), βIII-tubulin (Sigma-Aldrich, T8660), MF20 (DSHB, RRID:AB_2147781) were incubated overnight at 4 °C. The cells were then incubated with secondary antibodies: AlexaFluor 594 conjugated goat anti-mouse IgG, AlexaFluor 594 conjugated donkey anti-rabbit IgG (ThermoFisher). Samples without primary antibody were used as negative controls.

### 2.7 Hematopoietic differentiation

ICF2- and control-iPSCs were differentiated into hematopoietic progenitors using the STEMdiff Hematopoietic Kit (StemCell Technologies). Briefly, one day before starting the differentiation protocol, iPSCs at 70-80% confluency were treated with Cell Dissociation Buffer to obtain aggregates (100-200μm in diameter). A total of 100-150 clusters were transferred into a Geltrex-coated 6-well culture dish and cultured overnight. The following day the StemMACS iPS-Brew-XF was replaced with Basal medium with supplement A to induce iPSCs towards a mesoderm-like state. On day 2, half of the medium was changed. On day 3, Basal medium with supplement B replaced Medium A to promote differentiation into hematopoietic cells. Half medium changes were performed on days 5, 7 and 10. On day 12, floating and adherent cells were collected and prepared for FACS analyses.

### 2.8 Flow cytometry analyses

SSEA4 antigen was detected using a mouse primary antibody anti-SSEA4 (SCBT, sc-21704) and a secondary antibody conjugated with AlexaFluor 594. Cells were then analyzed by FACS. At the end of the hematopoietic differentiation protocol (Day 12), floating and adherent cells were incubated with a mix of antibodies. The following primary monoclonal antibodies (eBioscience) were used for cell surface antigen detection: anti-CD43 PE (12-0439-42), anti-CD45 FITC (11-0459-41) and anti-CD34 APC (17-0349-41). Isotype control was performed with monoclonal mouse IgG1 kappa Isotype Control PE (12-4614-81), FITC (11-4714-81), APC (17-4714-41). Live cells were gated based on FSC/SSC parameters. All phenotypes were analyzed with the FACSAria III system (BD Biosciences), acquiring 10000 events for samples. Data obtained were analyzed and elaborated by DIVA software (v8.0.1).

### 2.9 DNA methylation array analysis

Genomic DNA was extracted from iPSCs and blood using the Wizard Genomic DNA kit (Promega) following the manufacturer’s instructions. DNA was treated with sodium bisulfite and bisulfite-converted DNA was subjected to methylation profiling using Infinium EPIC-850K Array (Illumina). Fluorescence signal intensities were captured using Illumina HiScanSQ.

Using the “champ.load” function, the raw idat files of pYM-iPSCs, pYM-blood, control iPSCs and control blood samples (GSE237676; (26)) were imported to calculate the β-values with quality control and filtering options set as default and array type as “EPIC”. To normalize the methylation signal from Type I and II probes, the BMIQ normalization using the “champ.norm” function was applied with default parameters and array type as “EPIC”. Data derived from controls and ICF1-4 patients’ blood were kindly provided by Dr. Guillaume Velasco (10). The common probes between the arrays were considered using the combineArray function in minfi Bioconductor package v1.40.0. Using a linear model built with “limma” package, the differentially methylated CpGs were calculated with a mean β-values difference ≥|0.10| and adjusted P-value < 0.05.

To analyze the repeated regions, the coordinates from UCSC (http://hgdownload.cse.ucsc.edu/goldenpath/hg19/database/rmsk.txt.gz) were downloaded and the CpGs were annotated to the Satellite repeats (probe covering ≥10).

### 2.10 Statistical analysis

Data obtained from qPCR and flow cytometry experiments are presented as mean ± SD from three independent experiments. Statistical analyses were performed using the one-side two-sample Student’s t-test; (∗) P-value < 0.05, (∗∗) P-value < 0.01, (∗∗∗) P-value < 0.001, (∗∗∗∗) P-value < 0.0001. The statistical significance of the methylation differences between ICF2-iPSCs and the median of controls was calculated using a nonparametric two samples two-sided Wilcoxon test with BH-FDR correction. Statistical significance was calculated by comparing each clone of ICF2-iPSCs to the average of control-iPSC clones.

## 3 Results

### 3.1 Generation of ICF2-iPSCs from peripheral CD34^+^-blood cells of a patient carrying the p.Cys408Gly ZBTB24 pathogenic variant

To investigate the earliest events of molecular alterations in a ICF type 2 syndrome patient, we generated iPSCs from peripheral blood of the pYM patient carrying the homozygous missense mutation c.1222T>G (p.Cys408Gly) in exon 5 of *ZBTB24* (25). CD34^+^ cells were isolated from PBMCs and expanded (27). Reprogramming was induced through the nucleofection of episomal reprogramming vectors encoding for the pluripotency factors Oct4, SOX2, KFL4, L-Myc, Lin28 and mp53DD and EBNA1 proteins (28). A similar procedure was applied to generate iPSCs from peripheral blood of a healthy donor individual. Several clones were characterized and validated as bona fide iPSCs. The molecular validation of representative clones ICF2 pYM C6 and control C14 are shown in Figure 1A-C and Figure S1A-C, respectively. The chromosomal integrity of each clone was verified (Figures 1D and S1D). The ability of iPSC clones to spontaneously differentiate into the three embryonic layers was validated by immunofluorescence (Figures 1E and S1E). Loss of the episomal vectors during iPSC passaging was evaluated through a PCR assay specific for the exogenous sequences (Figure S1F).

**Figure 1.**
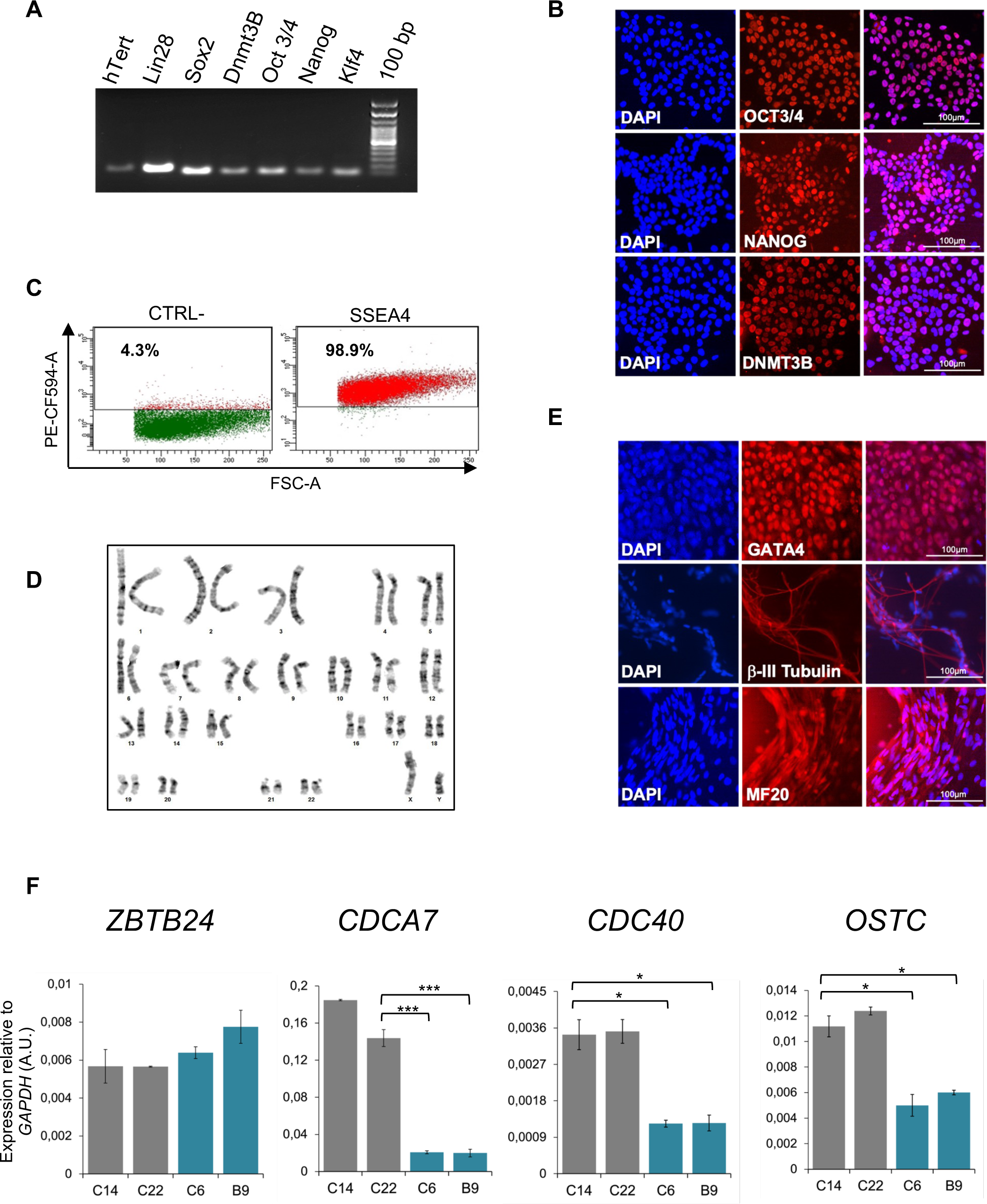
Molecular characterization of pYM-iPSCs. (**A**) Expression of pluripotency markers *hTERT, LIN28, SOX2, DNMT3B, OCT3/4*, *NANOG* and *KLF4* transcripts measured by RT-PCR in a representative clone of pYM-iPSCs (C6); (**B**) Expression of OCT3/4, NANOG and DNMT3B pluripotency markers measured by immunofluorescence in the pYM C6 clone. DAPI, nuclear staining. Bar: 100μm; (**C**) Dot plot showing the expression of the pluripotency marker SSEA4 in the pYM C6 clone compared to the negative control; (**D**) Karyotype analysis demonstrating chromosomal integrity of the pYM C6 clone; (**E**) Immunostaining of markers representing the three embryonic germ layers, GATA4 (endoderm), βIII-tubulin (ectoderm) and MF20 (mesoderm) expressed in the *in vitro* differentiated C6 clone. Bar: 100 μm; (**F**) *ZBTB24*, *CDCA7*, *CDC40* and *OSTC* transcripts level in control (C14 and C22) and pYM (C6 and B9) iPSC clones measured by qPCR. The relative expression compared to the expression of *GAPDH* is shown for each gene. qPCR data are presented as mean ± SD from independent triplicates. Statistical analyses were performed using a one-side two-sample Student’s t-test compared to control: (∗) P-value < 0.05, (∗∗) P-value < 0.005, (∗∗∗) P-value < 0.0005.

It has been reported that ZBTB24 directly binds the promoter region of its target genes *CDCA7, CDC40* and *OSTC,* and is required for their expression (20,22). The p.Cys408Gly variant does not alter ZBTB24 protein expression, but affects its ability to bind the target genes. Consistent with these results, ICF2- and control-iPSCs show comparable levels of the *ZBTB24* transcript, but reduced levels of *CDCA7, CDC40* and *OSTC* transcripts are clearly observed only in the ICF2-iPSCs (Figure 1F).

### 3.2 ICF2 pYM-iPSCs recapitulate the DNA methylation defects associated with ZBTB24 deficiency

All ICF syndrome subtypes are characterized by variable hypomethylation of pericentromeric satellite repeats. Additional loss of methylation at centromeric satellites is a distinctive trait of ICF2-4 patient subtypes, compared to ICF1 patients (10). To explore whether pYM-iPSCs show this phenotype, we performed whole genome DNA methylation analysis using the Infinium EPIC Array. We compared the DNA methylation levels of the ICF2 iPSC clones to those of two control iPSC clones. In particular, we analyzed DNA methylation levels of satellite repeats specifically positioned at centromeres (ALR/Alpha), at pericentromeric regions (GSAT, GSATX, GSATII, SST1) and those dispersed across the genome (BSR/Beta, SATR1, SATR2) with a coverage of ≥10 probes (Figure 2A). Consistent with the ICF2 phenotype (10), the majority of these repetitive sequences were significantly hypomethylated in both clones of pYM-iPSCs compared to the average of normal iPSCs.

**Figure 2.**
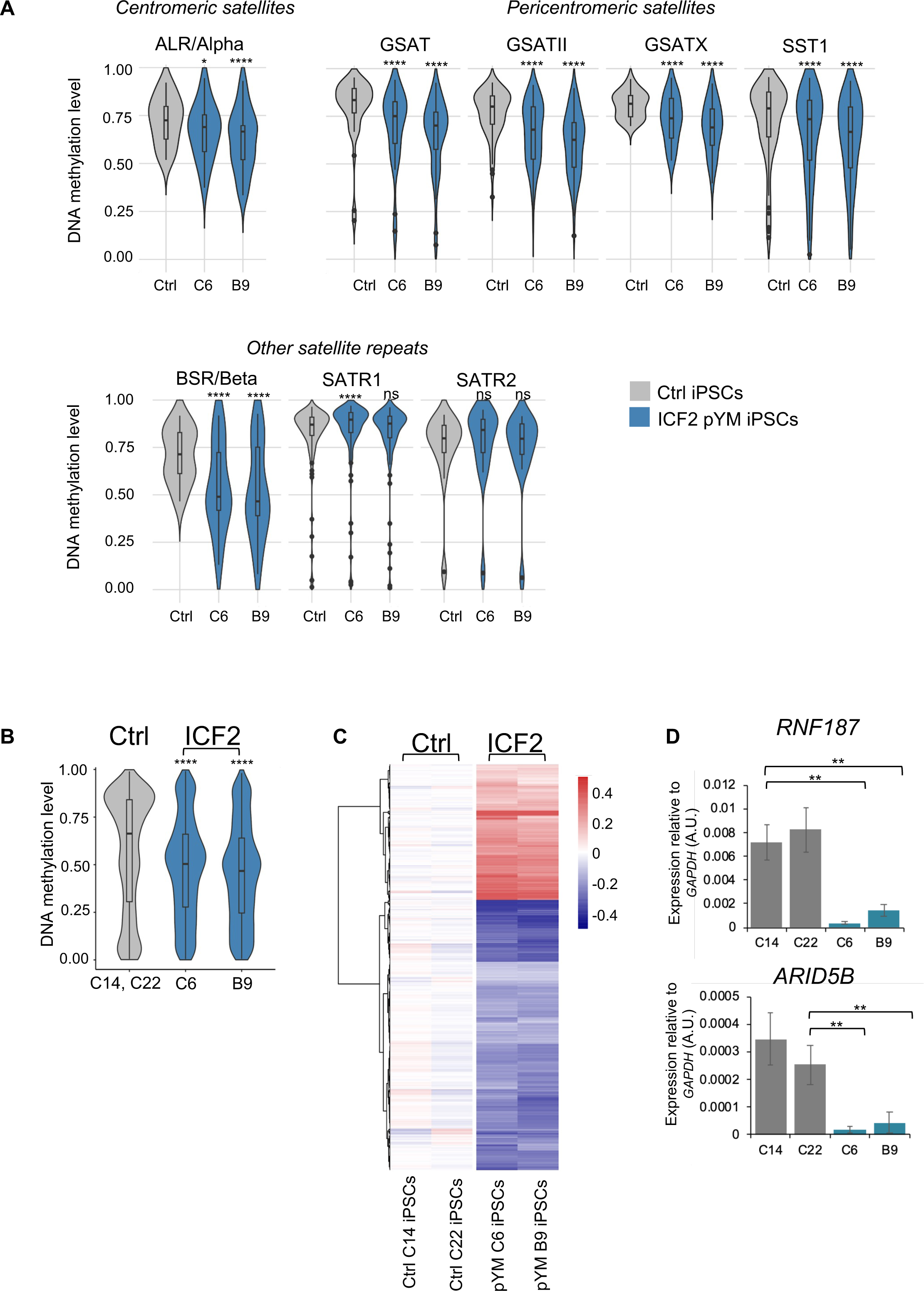
DNA methylation analysis in pYM-iPSCs. (**A**) Violin plot indicating the distribution of methylation values of Satellite repeats localized in centromeric (ALR/alpha), pericentromeric (GSAT, GSATII, GSATX, SST1) and other genomic regions (BSR/Beta, SATR1, SATR2) in averaged control-iPSCs (C14 and C22) and pYM-iPSCs (C6 and B9); (**B**) Violin plot showing the distribution of methylation values detected at DMPs in pYM-iPSCs compared to the average of methylation values in control-iPSCs. Boxes inside the violin are interquartile ranges with the median line. *P*-adjusted values represent the BH-corrected *p*-values obtained from a two-samples Wilcoxon test with two-sided alternatives; (**C**) Heatmap representing the value of methylation difference at 2753 DMPs in pYM-iPSCs compared to control. Individual sample methylation (β-values) is subtracted from the mean methylation (mean β-values) of the two controls. The blue and red represent the hypo- and hypermethylation level, respectively; (**D**) *RNF187* and *ARID5B* transcripts level in control and pYM-iPSC clones measured by qPCR. The relative expression compared to the expression of *GAPDH* is shown for each gene. qPCR data are presented as mean ± SD from independent triplicates. Statistical analyses were performed using a one-side two-sample Student’s t-test compared to control: (∗) P-value < 0.05, (∗∗) P-value < 0.005, (∗∗∗) P-value < 0.0005.

At the genome-wide level, we identified 2753 differentially methylated probes (DMPs; 960 linked genes) in pYM-iPSCs that overall showed significantly lower methylation levels than the averaged control-iPSCs (Figure 2B). While the majority of DMPs were hypomethylated in the patient iPSCs, 920 CpGs were hypermethylated when compared to control-iPSCs (Figure 2C and Table S2). Clusters of functionally related genes were enriched among the hypomethylated genes (Olfactory Receptors, Late Cornified Envelope, Defensin, Keratin-Associated Protein and Zinc-Finger proteins; Table S3). Genes involved in several biological processes were affected, including developmental pathways, synaptic signaling, nervous system process and natural killer cell mediated immunity (Table S3).

A subset of hypermethylated genes in ICF2-iPSCs (*ARID5B*, *RNF187, ARL11, COPB1, PLRG1*) were previously described as direct targets of ZBTB24 binding in normal lymphoblastoid cell lines (LCLs) and aberrantly silenced in patient LCLs (11). We thus tested the expression of these genes in control- and pYM-iPSCs. Interestingly, *ARID5B* and *RNF187* expression was significantly reduced in patient pYM-iPSCs, confirming that ZBTB24 LOF is associated with promoter hypermethylation and silencing of these genes also in iPSCs (Figure 2D).

The biological processes of genes affected by DNA methylation alterations were similar to those that characterize ZBTB24 deficiency in ICF2 patients’ blood. Several of them were related to nervous system functions, consistent with the neurodevelopmental delay characteristic of ICF2 patients (16,25). The DNA methylation profile of pYM blood was compared to that of the other known ICF2 patients, taking advantage of previously published methylation-array based data (10). We found that pYM blood methylation clearly clustered together with those of ICF2-4 patients and was distinct from the ICF1 and ICFX subgroups, which are known to be closer to each other (Figure S2).

Next, we identified the specific fraction of DMPs (1612; 552 linked genes) in pYM-iPSCs that were originally present in the peripheral blood of the patient, by comparing the methylation array results in pYM iPSCs and whole blood (Figure 3A and Table S4). Among the hypomethylated regions, we detected the olfactory receptors cluster and several other gene clusters (Figure 3B and Table S5). Notably, *ARID5B* and *RNF187* were hypermethylated in both pYM iPSCs and blood samples (Figure 3C).

**Figure 3.**
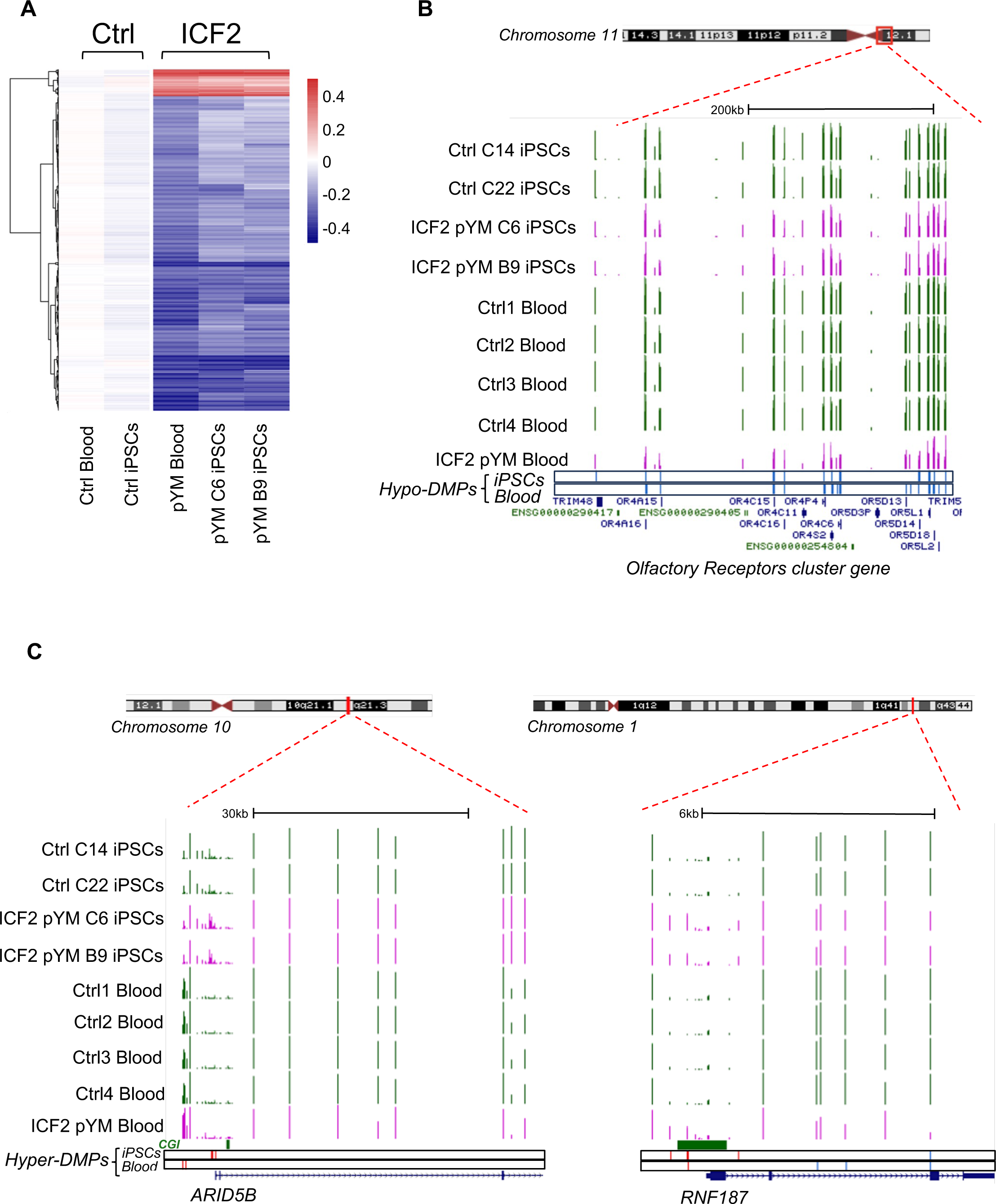
DNA methylation analysis in pYM blood and iPSCs. (**A**) Heatmap representing the value of methylation difference at 1612 DMPs that are detected in both pYM blood and iPSC clones. Individual sample methylation (β-values) in ICF2 patient samples is subtracted from the mean methylation (mean β-values) of blood and iPSC control samples; (**B**) Genome browser view of the centromeric and pericentromeric region at chromosome 11 including the clustered Olfactory Receptor genes. The coverage tracks from the top to the bottom display the DNA methylation levels shown as β-values for control and pYM-iPSC clones and for control and pYM blood samples. The underneath blue bars represent the hypo-DMPs in patient iPSCs and blood, respectively; (**C**) Genome browser view of the *ARID5B* and *RNF187* loci. The underneath red and blue bars represent the hyper- and hypo-DMPs in patient iPSCs and blood. CpG islands and the exon/intron boundaries are also indicated.

Overall, we demonstrated that the iPSCs generated from pYM patient CD34^+^ cells recapitulate the ICF2-specific methylation defects, rendering this platform as suitable for the study of pathogenetic mechanisms in ICF2.

### 3.2 ICF2 pYM-iPSCs differentiate towards hematopoietic progenitors with reduced efficiency

To investigate the earliest phenotypic alterations that characterize ZBTB24 LOF, we directed pYM- and control-iPSCs to differentiate towards hematopoietic progenitor cells. We utilized a monolayer based feeder-free protocol for hematopoietic differentiation (29) that enables the generation of CD34, CD43 and CD45 positive cells. We estimated cell survival for control and pYM derivatives by the end of the differentiation process. The percentage of live cells was respectively 65.1% for control and 49.3% for pYM derivatives, indicating a different survival rate during the differentiation (Figure 4A). The lower vitality of the pYM derivatives was confirmed in three independent experiments (Figure 4B). Living cells were further evaluated for the expression of two hematopoietic markers, CD43, the earliest marker of human hematopoietic commitment, and CD45, which is the pan-hematopoietic marker gradually acquired during hematopoietic differentiation. Through this analysis we visualized two distinct populations both in pYM and control cells (Figure 4C). The most abundant population is positive for CD43 marker expression and presents different levels of CD45 marker (identified as CD43^+^/CD45^+/-^). The less abundant population is identified as CD43^-^/CD45^-^ and is characterized by the absence of both hematopoietic markers. These experiments highlighted different ratios between “hematopoietic” and “non-hematopoietic” populations derived from ICF2- and control-iPSCs. While the CD43^+^/CD45^+/-^ cells were slightly decreased in ICF2 derivative cells compared to control, the double negative cells were significantly increased in ICF2 cells (Figure 4D).

**Figure 4.**
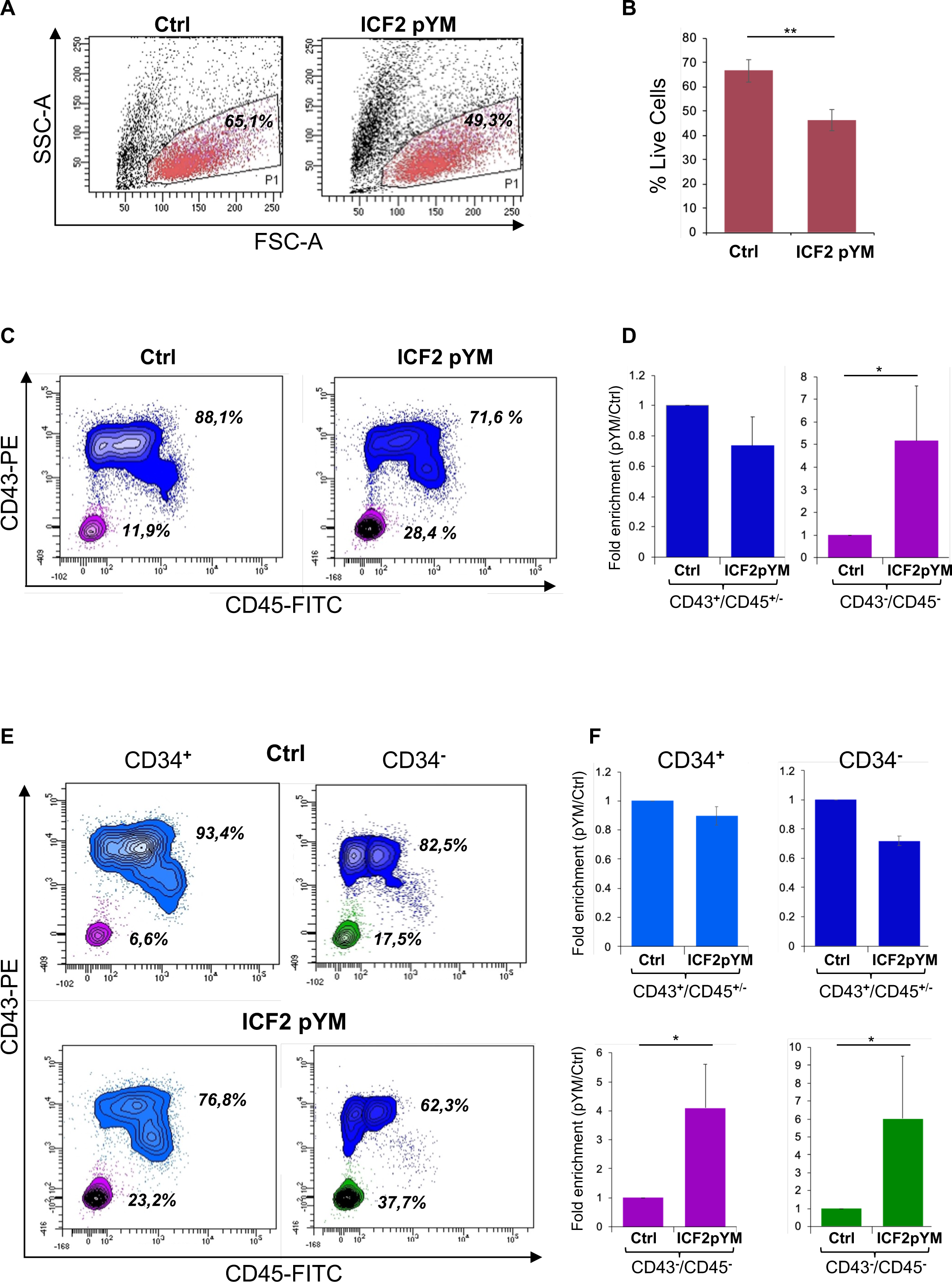
Differentiation of pYM-iPSCs toward hematopoietic progenitors. (**A**) FSC-A/SSC- A morphological dot plot of living cells derived from hematopoietic differentiation of control- and pYM-iPSCs. The percentages are relative to the representative experiment; (**B**) Percentages of live cells derived from control and pYM-iPSCs in three independent experiments presented as mean ± SD; (**C**) Contour plot showing the expression of CD43 and CD45 in the total live cell population. The percentages are relative to the representative experiment; (**D**) Fold enrichment of CD43^+^/CD45^+/-^ and CD43^-^/CD45^-^ of pYM derivatives compared to control cells in three independent experiments presented as mean ± SD; (**E**) Contour plot showing the expression of CD43 and CD45 evaluated on the selected CD34^+^ and CD34^-^ cells. The percentages are relative to the representative experiment. (**F**) Fold enrichment of CD43^+^/CD45^+/-^ and CD43^-^/CD45^-^ cells evaluated on the selected CD34^+^ and CD34^-^ cells in three independent experiments presented as mean ± SD. Statistical analysis was performed using a one-side two-sample Student’s t-test compared to control: (∗) P-value < 0.05, (∗∗) P-value < 0.005, (∗∗∗) P-value < 0.0005.

As CD34 is also expressed in non-hematopoietic progenitors, we investigated the expression of this marker in living cells derived from the differentiation of ICF2- and control-iPSCs. The analysis showed comparable levels of CD34 expression in control and ICF2 cells (Figure S3A,B). We then examined the co-expression of CD43 and CD45 markers in the CD34 positive and negative cells. The analysis revealed a marked increase of triple negative CD34^-^/CD43^-^/CD45^-^ and of CD34^+^/CD43^-^/CD45^-^ cells in ICF2 derivative cells compared to control cells (Figure 4E, F). Overall, our results demonstrate that the pYM-iPSCs generated hematopoietic progenitors with lower efficiency than control cells. Further, they showed an increased percentage of non-hematopoietic CD43^-^/CD45^-^ cells.

## 4 Discussion

ICF syndrome is an ultra-rare disease with genetic and clinical heterogeneity. DNA methylation is the pivotal molecular hallmark that exerts pleiotropic effects leading to a wide range of impairments, which result in a multisystemic syndrome. Disruptions in the epigenome are increasingly being recognized as a common pathogenic mechanism underlying rare Inborn Errors of Immunity (30). DNMT3B is the only gene among those involved in the ICF syndrome pathogenesis that is clearly associated with DNA methylation homeostasis (31). About 40 ICF2 patients carrying *ZBTB24* variants have been described (16). Immunological features of ICF2 patients are extremely variable ranging from very mild to severe immunodeficiency (14,15,25,32,33). Moreover, a progressive course of immune dysfunction has been described in some ICF2 cases (4). Clinical heterogeneity and lack of clear genotype/phenotype correlation among the patients despite the common genetic background point to the relevance of epigenetic modifications in the disease pathogenesis.

Mouse models carrying homozygous mutations in the BTB domain show embryonic lethality, supporting the notion that molecular alterations arise during early development (22). Overall, the studies in humans (34), mice (22) and zebrafish (35) suggest that ZBTB24 is vital for normal progression through early developmental phases. Notably, reduced levels of *ZBTB24* transcripts were recently observed in decidua and placental villi derived from women with recurrent spontaneous abortions (RSA). Furthermore, *ZBTB24* silencing in cells derived from human trophoblasts modifies the expression of E-cadherin by altering DNA methylation levels at the promoter region. This evidence supports the notion that epigenetic changes associated with ZBTB24 activity may interfere with early embryo development (34).

Here, we generated the first iPSC-based model for ICF syndrome type 2. We reprogrammed CD34^+^ cells from a patient homozygous for p.Cys408Gly. This human cellular model recapitulated the transcriptional and epigenetic abnormalities associated with ZBTB24 deficiency. We analyzed the transcript level of the *CDCA7, CDC40* and *OSTC* genes that are direct targets of the ZBTB24 transcription factor (22). Consistent with the results of the *Zbtb24*-deficient mESCs, these three genes showed significantly reduced expression in pYM-iPSCs. Altogether, these data suggest that ZBTB24 is a transcriptional activator also in iPCSs. Moreover, we found that the typical ICF2-specific methylation defects at centromeric and pericentromeric satellite repeats are recapitulated in patient-derived iPSC clones, with a consistent overlap among different clones.

The ability of ZBTB24 to bind centromeric satellite DNA and play a critical role in establishing the correct DNA methylation pattern at these regions, through the recruitment of DNMT3B, has been recently proposed in mESCs (11,36). The levels of minor satellite transcripts at mouse centromeres were significantly higher in *Zbtb24*mt/mt mESCs compared to wild type mESCs. In ICF2 patient LCLs, ZBTB24 LOF causes higher levels of ALR/alpha satellite and satellite 2 transcription (11,36) and associated chromosomal instability (37), suggesting that this protein acts as a guardian of centromeric integrity (11). Our data in pYM-iPSCs support the notion that ZBTB24 is an early factor required for the correct centromeric DNA methylation. We previously generated ICF1 patient-derived iPSCs that are deficient for DNMT3B (38–40). Comparative mechanistic studies in ICF2 and ICF1 iPSCs will provide insights into the functional interaction between the two proteins and their regulatory pathways.

DNA methylation was similarly perturbed at several gene clusters in the ICF2-patient iPSCs. These included Olfactory Receptors, Keratin-associated proteins, Defensins and Protocadherins that were similarly hypomethylated in the patient’s blood. These genes have been typically found hypomethylated in all ICF2 patient blood and LCLs (10). In pYM-iPSCs we also detected hypomethylation of genes involved in neural development and immune function. Of note, genes playing roles in natural killer cell mediated immunity also showed an altered DNA methylation pattern, consistent with the immunodeficiency phenotype. Hence, our findings suggest that DNA methylation of these genes might be primed during early stages of disease development before their activation in the proper cellular context.

DNA hypermethylation patterns were also detected in ZBTB24-deficient iPSCs and numerous genes acquired promoter methylation. Among them, *ARID5B* and *RNF187* showed aberrant gene silencing in these cells. These genes are direct targets of ZBTB24 and were transcriptionally downregulated in ZBTB24-deficient LCLs and in ZBTB24-mutant mESCs (11,14). Several studies indicate that ZBTB24 binding protects the promoter of these genes from DNMT3B-mediated methylation and gene silencing (41). Whether ZBTB24 directly influences DNA methylation at gene promoters is still unclear. Genome-wide transcription studies combined with the ZBTB24 binding profile will further clarify the direct and indirect consequences of ZBTB24 deficiency on regulation of gene expression.

The link between ZBTB24 impairment and clinical phenotype is still elusive. ZBTB proteins are involved in hematopoietic development and lineage commitment, but the specific contribution of ZBTB24 is unknown. There is evidence that it is required for normal B and T cell proliferation and function, regulating the expression of lineage-specific factors (42–45) Notably, in patient-derived iPSCs we observed a significant downregulation of *CDCA7* that is a Notch transcriptional target involved in hematopoietic stem cell emergence (46). Therefore, it is plausible that the transcriptional irregularities due to epigenetic abnormalities in pYM-iPSCs emerge during the specification of hematopoietic stem and progenitor cells. The ICF2-iPSC model allowed us to investigate the role of ZBTB24 in this process. Interestingly, patient-derived iPSCs showed a reduced vitality during the differentiation process, suggesting a potential activation of apoptosis. The ZBTB24-CDCA7 axis has been implicated in the regulation of cell-cycle progression and proliferation (43,45) and its impairment can be associated with the phenotype that we observed in ICF2 cells.

Hematopoietic CD34/CD43/CD45 positive progenitors were generated with lower efficiency, while the non-hematopoietic fraction increased in the ICF2 derivative cells. This latter subpopulation might include a higher proportion of endothelial or mesenchymal progenitors (29). Future analyses addressing the detailed composition of the cellular subpopulations that emerged in patient iPSC derivatives will contribute to clarification of the phenotypic defects occurring during hematopoietic differentiation.

Our findings demonstrate that the ICF2-iPSC model is highly relevant for studying the role of the ZBTB24 transcription factor in the DNA methylation homeostasis. We anticipate that this novel platform will provide a versatile tool to shed light on the complex cascades of early molecular events linking *ZBTB24* deficiency to the clinical phenotype of ICF type 2 syndrome.

## Data availability statement

Illumina EPIC array raw and processed data are deposited in the GEO repository under accession GSE262957.

## Author Contributions

Conceptualization, MRM and MS; Investigation VL, FC, SB, SA, BM; Validation VL, SB, SA, BM, SF, FDR, RG; Resources, GG, CP; Visualization VL, FC, SB; Formal analysis, FC, LP; Writing original draft preparation, VL, FC, MRM, MS; Writing, Review and editing VL, FC, SV, SA, BM, SF, LP, RG, FDR, GG, CP, AR, MRM, MS; Supervision, MRM, MS; Funding acquisition, FDR, AR, MRM, MS.

## Acknowledgment

We acknowledge Vincenzo Mercadante (FACS Facility), Rosarita Tatè and Salvatore Arbucci (Integrated Microscope Facility IGB-IM) for their valuable help with the flow cytometry and imaging experiments. We thank Domenico Marano and Annalisa Fico for their help with the immunofluorescence experiments. We are grateful to Guillaume Velasco and Claire Francastel for sharing the Illumina 450K methylation data. We also thank Imma Di Biase (Merigen Diagnostic) for the contribution to the karyotype analysis, and Lorenzo Chiariotti and Mariella Cuomo for their support in the DNA methylation array experiments. Many thanks to Sara Selig for helpful advice on the experiments and comments on the manuscript.

