## Supplementary Figures for "A novel iPSC-based model of ICF syndrome subtype 2 recapitulates the molecular phenotype of ZBTB24 deficiency"

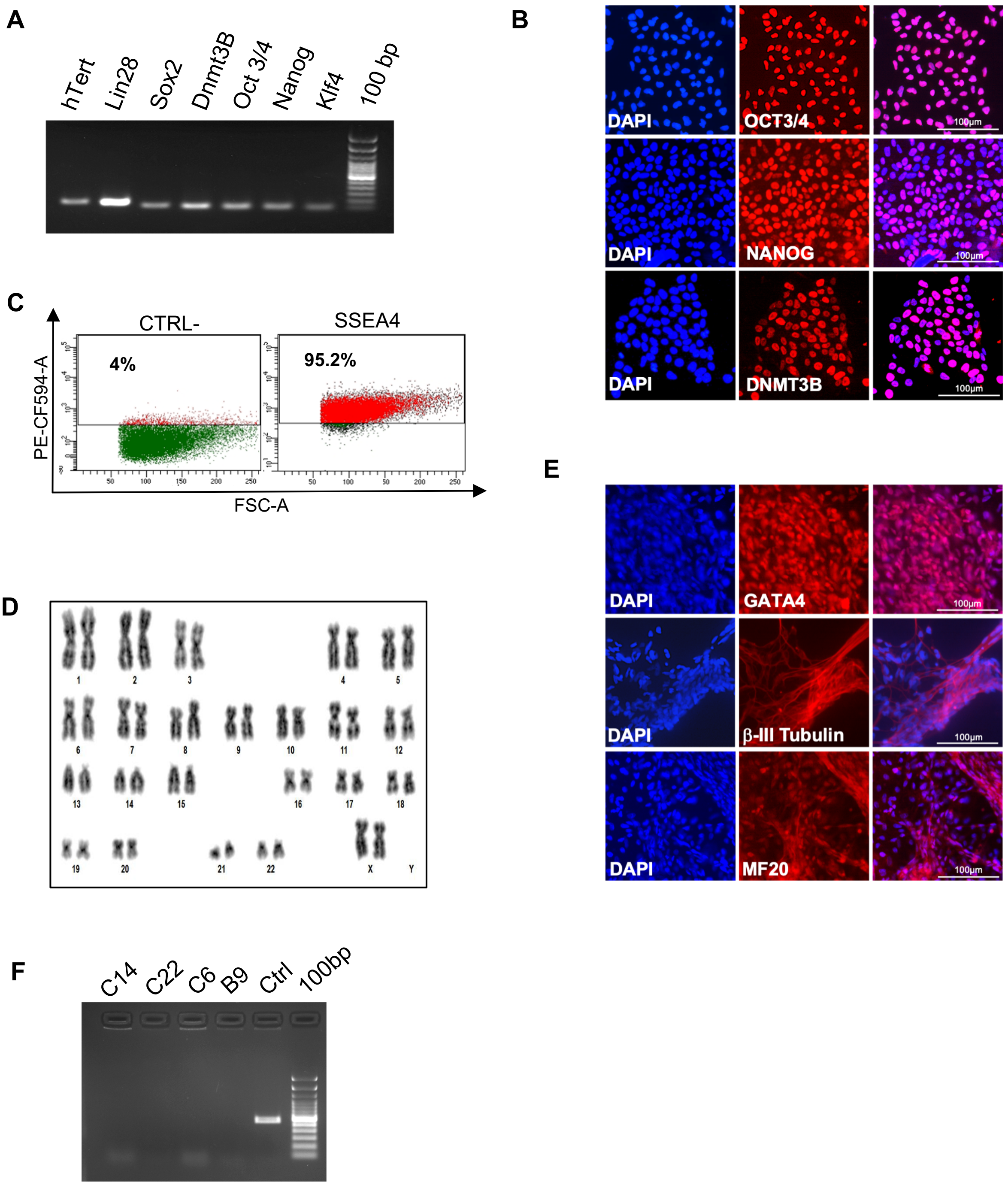


**Supplementary Figure S1.** Molecular characterization of control-iPSCs. (**A**) Expression of pluripotency marker *hTERT, LIN28, SOX2, DNMT3B, OCT3/4*, *NANOG* and *KLF4* transcripts measured by RT-PCR in a representative clone of control-iPSCs (C14); (**B**) Expression of OCT3/4, NANOG and DNMT3B pluripotency markers by immunofluorescence in the C14 clone. DAPI, nuclear staining. Bar: 100 µm; (**C**) Dot plot showing the expression in the C14 clone of the pluripotency marker SSEA4 compared to the negative control; (**D**) Karyotype analysis demonstrating the chromosomal integrity of the C14 clone; (**E**) Immunostaining of markers representing the three embryonic germ layers, GATA4 (endoderm), βIII-tubulin (ectoderm) and MF20 (mesoderm) expressed by the *in vitro* differentiated C14 clone. Bar: 100 µm; (**F**) Loss of the transfected episomal vectors in control- and pYM-iPSCs evaluated by PCR.

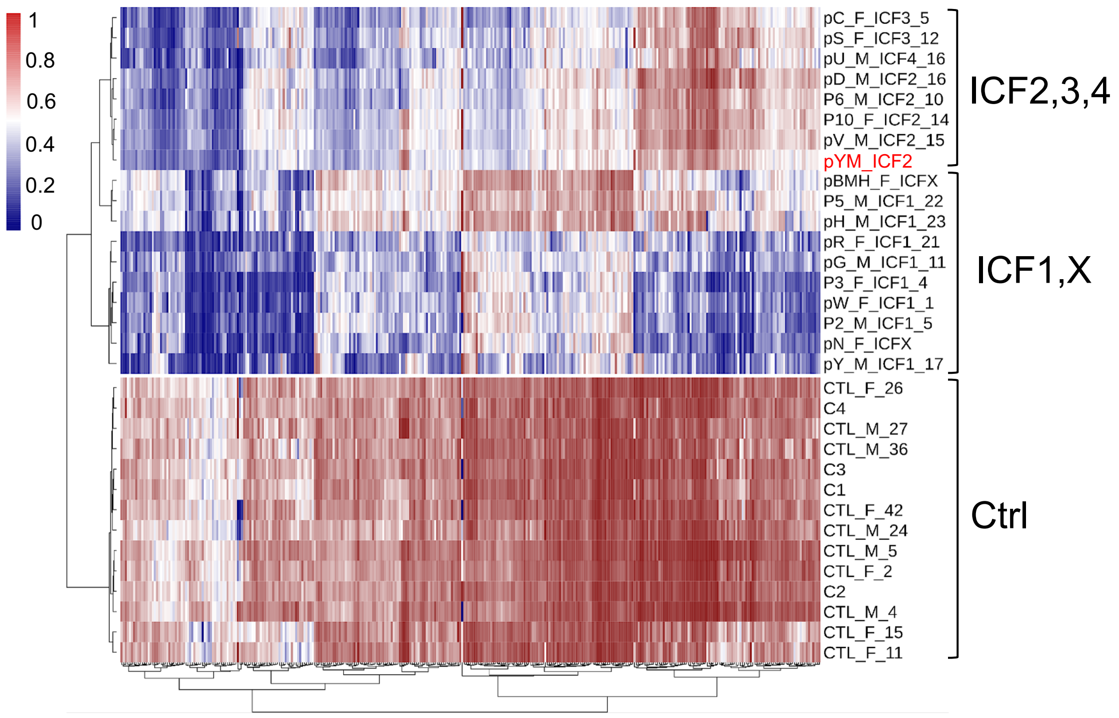


**Supplementary Figure S2.** Comparative analysis of DNA methylation profiles in ICF patient blood samples. Heatmap representing the value of methylation levels in the pYM sample compared to control, ICF1-4 and ICFX patient samples (10).

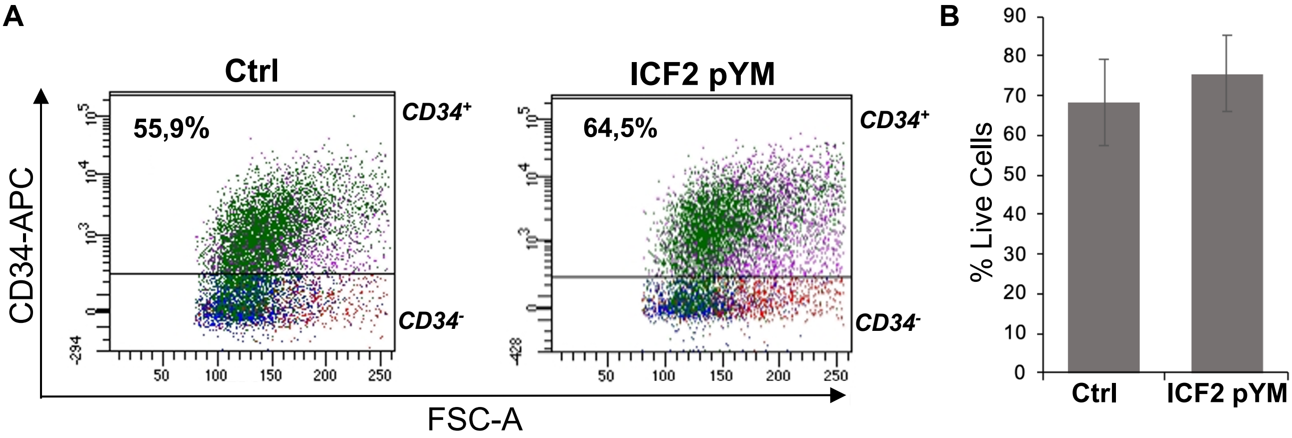


**Supplementary Figure S3.** Expression of CD34 marker in control and pYM differentiation derivatives. (**A**) Dot plot showing the expression of the surface marker CD34 in control and pYM cells generated from hematopoietic differentiation. The number of positive cells for the marker is expressed as the percentage of live cells detected in this representative experiment; (**B**) Percentage of positive live cells derived from control- and pYM-iPSCs in three independent experiments presented as mean ± SD.
